# An Integrated Multiphoton Imaging Workflow for Quantitative Analysis of Aortic Tissue Microstructure

**DOI:** 10.64898/2026.01.25.701601

**Authors:** Mirza Muhammad Junaid Baig, Ana I. Vargas, Turner Jennings, Rouzbeh Amini, Chiara Bellini

## Abstract

Quantitative, reproducible characterization of aortic microstructure is essential for advancing vascular biomechanics and mechanobiology. To address this need, we present a comprehensive image-analysis workflow that extracts quantitative descriptors of tissue microstructure from multiphoton microscopy stacks of the murine thoracic aorta. Channel-specific signals are acquired for fibrillar collagen (second harmonic generation), elastin (two-photon autofluorescence), and cell nuclei (two-photon excited fluorescence). Following reorientation into the *XZ* plane, individual elastic lamellae are traced to quantify lamellar thickness and interlamellar spacing using circle-based geometry (Taubin fitting). After correction for vessel wall curvature via a cylindrical transformation, segmented nuclei are assigned to medial or adventitial compartments based on visual estimates of adventitial volume fraction, and nuclear morphology is characterized via ellipsoidal fitting in terms of nuclear aspect ratio and major-axis orientation. Collagen organization is resolved in *XY* sections by extracting fiber centerlines to quantify straightness and amplitude; traces from serial sections are then combined to reconstruct the three-dimensional collagen network and estimate porosity and linear fiber density, while fiber orientation distributions are derived from principal component analysis–based angles and fit using a von Mises mixture model. Finally, collagen and elastin volume fractions are computed via a two-stage fixed-threshold approach calibrated on a balanced training subset. Overall, this modular and robust workflow provides an integrated framework for studying aortic wall remodeling across physiological and pathological processes.

**Non-Technical Summary:** As the main blood vessel in our body, the aorta needs to be both strong and flexible. This balance comes from three main parts: elastic layers that allow the aorta to stretch, strong fibers that prevent tearing, and cells that sense and respond to changes in blood pressure and other signals. When any of these components are altered, the aorta may stiffen or weaken, which can interfere with normal blood flow. In this study, we developed a clear and consistent way to measure the structure of the aortic wall using microscope images. The approach examines how thick the elastic layers are and how far apart they lie, the size and orientation of cell centers, and how straight or wavy structural fibers appear. It also estimates how much of each component is present in the aortic wall. Because the same steps are applied each time, results can be fairly compared across different conditions. Overall, this tool transforms detailed images into simple measurements, helping scientists understand how the aorta changes in health and disease.

## 1 Introduction

The microstructural organization of the extracellular matrix (ECM), alongside the spatial arrangement of resident cells, confers mechanical integrity to soft biological tissues and underpins adaptive remodeling in support of physiological function [1, 2, 3, 4]. Elastic arteries, including the aorta, exhibit pronounced structure-function coupling, thus quantitative analyses of tissue microstructure are integral to vascular biomechanics and mechanobiology [5, 6, 7]. Such approaches translate imaging data into descriptors with direct relevance to tissue-level mechanical behavior [8, 9, 10, 11, 12], suitable for informing mechanistic hypotheses and constitutive modeling [13, 14]. On this basis, we sought to extract microstructural metrics from multiphoton image stacks of murine aorta in a reproducible manner, while retaining direct relevance to the mechanical contributions of individual wall compartments. Because collagen and elastin play distinct and complementary mechanical roles in the aortic wall, we first focused on these key structural components of the ECM [6]. Elastin forms lamellar units within the aortic media that dominate the mechanical response at low strain and provide elastic recoil over repeated loading cycles [15, 16, 17, 18, 19, 5]. Developmental and age-related changes in lamellar thickness, interlamellar spacing, and structural integrity alter wall mechanics, thus incorporating lamellar geometry into constitutive models improves low-strain predictions and more accurately captures load sharing between microstructural constituents [20, 21, 22, 23, 24, 25, 26, 27]. With increasing strain, collagen fibers dispersed throughout the aortic wall are progressively recruited to bear load [18, 20, 21, 22]. Collagen architecture, including orientation distribution, dispersion, and recruitment behavior, drives anisotropy and directional stiffness [28, 29, 30, 31, 32, 33]. Accordingly, multiscale modeling and direct-fiber simulations show that fiber organization gives rise to heterogeneous stress and strain fields, which shape macroscopic behavior and influence damage initiation [34, 35, 36, 37, 38, 39, 40]. Taken together, these observations underscore the need for measurements that resolve the microstructure of aortic ECM constituents across the physiological strain range.

Cells such as those residing in the vasculature dynamically sense mechanical cues in their environment and respond by modulating the microstructure of the surrounding ECM. In this process, the cell nucleus acts as a mechanosensitive, load-bearing organelle whose shape and orientation reflect the local strain pattern [41, 42, 43]. Nuclear morphology and deformation are associated with gene-expression programs and spatial strain fields in planar tissues, including those within the cardiovascular system [44, 45]. When analyzed alongside collagen and elastin architecture, nuclear metrics are thus expected to provide complementary insight into spatial gradients in mechanical signals across aortic wall compartments and the remodeling responses that follow altered loading [46, 47, 48, 49, 50]. This pairing bridges cell-scale observations with tissue-scale behavior without requiring invasive manipulation of the wall.

Multiphoton microscopy provides label-specific, depth-resolved readouts that are well suited for quantifying the spatial organization and distribution of collagen, elastin, and cell nuclei within intact aortic tissues. Second harmonic generation reports fibrillar collagen, while two-photon–excited fluorescence captures elastin and fluorophore-labeled nuclei, together enabling constituent-specific imaging across the aortic wall [51, 52, 53, 54]. Prior studies have used multiphoton and confocal imaging to delineate the layered architecture of the aortic wall and to characterize the three-dimensional (3D) structure of lamellar units within the media [55, 9, 56, 57]. Despite these advances, there remains a need for a comprehensive analytical framework that operates on raw image stacks, incorporates appropriate normalization and correction steps, accounts for curvature in resolving wall compartments and aggregating nuclear features, and yields collagen and elastin metrics pertinent to tissue mechanics [58]. Establishing such a workflow would reduce analyst dependence, facilitate direct comparison among experimental groups and conditions, and improve consistency of results across laboratories.

To construct this framework, two complementary strategies for quantifying collagen organization are available. Texture-based methods, including structure tensor analysis and wavelet or Riesz approaches, return orientation distributions and dispersion indices without explicitly resolving individual fiber centerlines; representative implementations include tools such as OrientationJ and FibrilTool [59, 60, 9, 61]. However, when applications require detailed fiber-level geometric features, including length, straightness or waviness, and curvature, explicit tracing approaches are preferred. Among these, Curvelet Transform Fiber Extraction (CT-FIRE) enhances elongated structures using a curvelet transform and segments individual fibers to extract centerlines and associated morphological metrics [62, 63]. Open-source alternatives such as FiberApp provide related capabilities for the analysis of filamentous networks [64]. Orientation histograms derived from these methods are commonly parameterized using von Mises mixture models comprising an isotropic background and a specified number of anisotropic fiber families. Such formulations map naturally to microstructural and microstructurally motivated constitutive laws for biological soft tissues that incorporate preferred directions and alignment parameters [65, 66, 67]. Together, texture-based measures and fiber-level descriptors have been leveraged to support comparative studies and constitutive model calibration in aortic tissues [57, 56].

Irrespective of the specific quantification strategy, standard image-processing considerations must be addressed to ensure reliable measurements. These include channel separation and correction for spectral cross-talk to minimize bias in constituent-specific metrics, compensation for depth-dependent attenuation in signal intensity, and conservative wall masking to define appropriate analysis domains [58, 63, 68]. In addition, transparent reporting of normalization choices, threshold values, and exclusion criteria facilitates sensitivity analyses to evaluate the robustness of the resulting measurements and strengthens the reliability of cross-sample comparisons [69, 70, 71, 72].

Building upon these considerations, we sought to develop and validate an integrated, curvature-aware analytical pipeline that uses multiphoton images of pressurized and axially extended murine aortic samples to (i) quantify elastic lamellar thickness and interlamellar spacing; (ii) assign cell nuclei to the appropriate wall compartment within a flattened coordinate system and derive principal nuclear orientation and shape; and (iii) extract collagen fiber features, including straightness, amplitude, porosity, and orientation distributions, to inform mixture-model parametrization. The purpose of this workflow is to extract constituent-level features that relate tissue microstructure to macroscopic mechanical behavior in a straightforward and reproducible manner. Together, these elements provide a consistent basis for comparing tissue microstructure across biological contexts and for informing constitutive models that account for geometry and load-bearing patterns [73, 74, 75, 76, 77, 78, 79, 80, 81].

## 2 Methods

### 2.1. Tissue Specimen Preparation

Multiphoton imaging data were obtained from segments of the descending thoracic aorta harvested from female C57BL/6 mice (10 *±* 1 weeks of age; n = 5). All experimental procedures adhered to National Institute of Health guidelines and were approved by the Northeastern University Institutional Animal Care and Use Committee (IACUC). Following excision, samples were cleaned to remove excess fat and perivascular tissue, and lateral branches were ligated with 9-0 nylon sutures. Specimens were incubated in SYTO™ Red nucleic acid stain (2 *μ*M; Invitrogen, Waltham, MA) for 2 h to visualize cell nuclei [68], then mounted on a custom-built pressurization device and stretched to approximate in vivo axial length based on established measurements in companion studies [82, 83].

### 2.2. Multiphoton Imaging Data Acquisition

3D multiphoton image stacks of the thoracic aorta were acquired to analyze elastin and collagen architecture, alongside nuclear morphology, under axially isometric and circumferentially isobaric conditions. Imaging was performed on a Zeiss LSM 880 NLO microscope with a W Plan-Apochromat 20 *× /*1.0 DIC (UV) VIS-IR M27 water-immersion objective and a tunable titanium–sapphire laser for excitation. The excitation wavelength was set to 843 nm for two-photon excitation. Emission signals were collected in separate spectral windows to isolate each constituent: 390-425 nm for second-harmonic generation (SHG) from fibrillar collagen, 500-550 nm for elastin two-photon autofluorescence (TPEF-AF), and *>* 550 nm for two-photon–excited fluorescence (TPEF) from SYTO-stained cell nuclei. Laser power was adjusted for each acquisition to prevent detector saturation while maintaining adequate image contrast. Representative *XY* optical sections from the elastin, nuclei, and collagen channels are shown in Fig. 1a.

**Figure 1.**
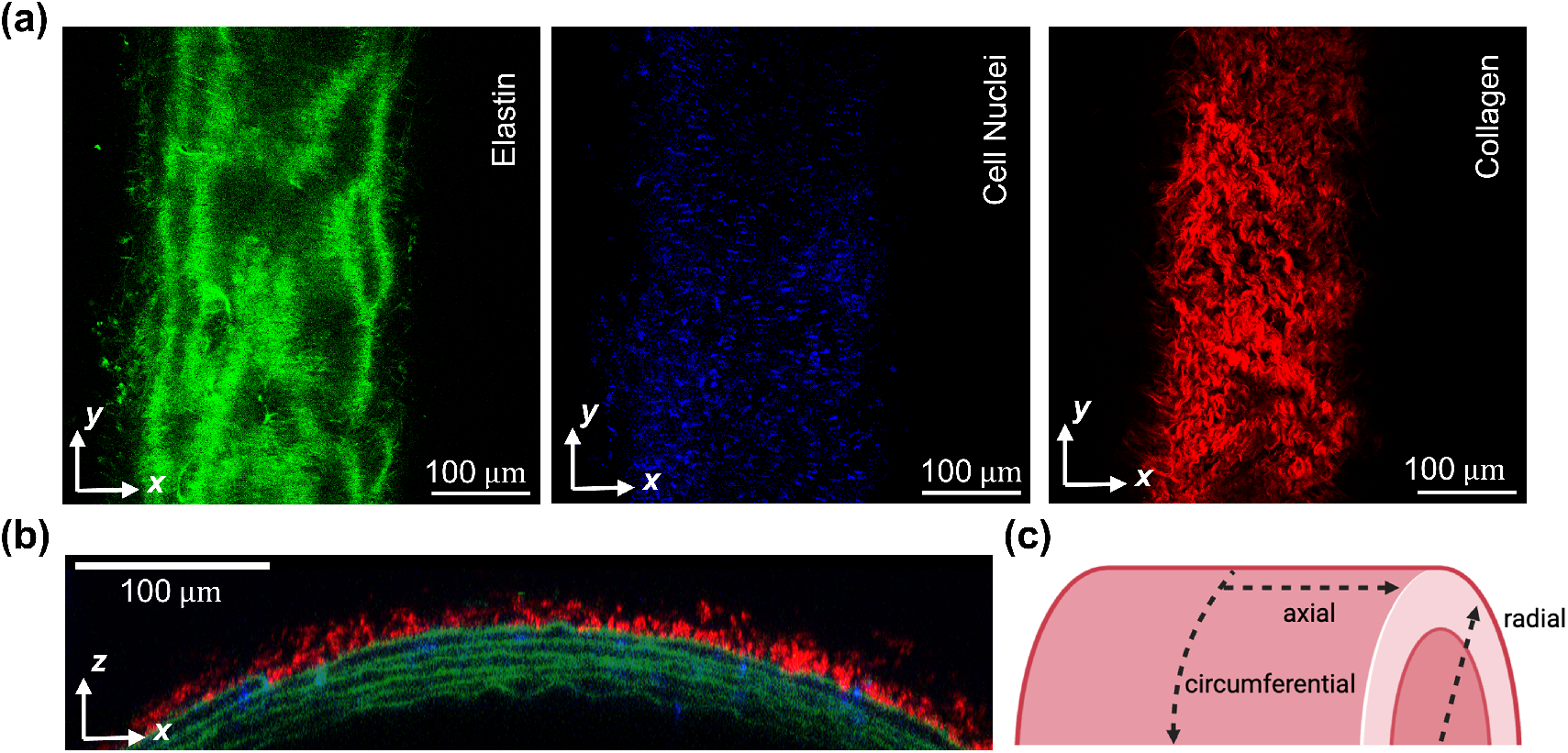
Multiphoton images and coordinate conventions. (a) Representative views of the three co-acquired channels: elastin (two-photon autofluorescence, green), cell nuclei (two-photon SYTO fluorescence, blue), and collagen (second harmonic generation, red). Images show individual *XY* optical sections from a representative 3D image stack. (b) Representative cross-sectional view of co-registered multiphoton signals illustrating the layered architecture of the aortic wall. Image combines individual co-registered *XZ* optical sections from a representative 3D image stack. (c) Schematics of the aortic wall visualizing the main anatomical directions: radial (through the wall thickness), circumferential (tangent to the vessel wall), and axial (parallel to the vessel long axis).

To visualize the layered wall architecture, co-registered *XZ* cross-sectional views were generated from each image stack (Fig. 1b). The radial, circumferential, and axial directions used throughout the analysis were defined relative to aortic wall anatomy, as illustrated schematically in Fig. 1c.

Each imaged stack covered a 500 *×* 500 *μ*m field of view with an *XY* in-plane resolution of 0.47 *×* 0.47 *μ*m per pixel. The axial step size along *Z* varied across samples according to acquisition settings. All subsequent image processing and quantitative analyses were performed in MATLAB R2024b.

### 2.3. Elastin Autofluorescence Image Analysis

Elastic lamellar architecture was resolved using the two-photon autofluorescent signal from elastin acquired in multiphoton *Z*-stacks of the aortic wall. Image stacks were reoriented from the native circumferential-axial (*XY*) plane into the circumferential-radial (*XZ*) view to visualize elastic lamellae within the medial compartment. The *XZ* projection of the elastin channel was binarized, and a central region of interest (ROI) was manually selected to avoid peripheral distortions. Within this ROI, individual lamellae were manually traced, and the corresponding boundary coordinates were extracted for geometric analysis (Fig. 2a). Lamellar thickness was quantified by fitting circles to the inner and outer boundaries of each traced lamella using the Taubin circle-fitting algorithm [74]. Radial thickness was then computed as the difference between the fitted radii, T_*n*_ = *R*_o,*n*_ − *R*_i,*n*_, where *R*_o,*n*_ and *R*_i,*n*_ are the radii of the outer and inner boundaries of the *n*-th lamella.

**Figure 2.**
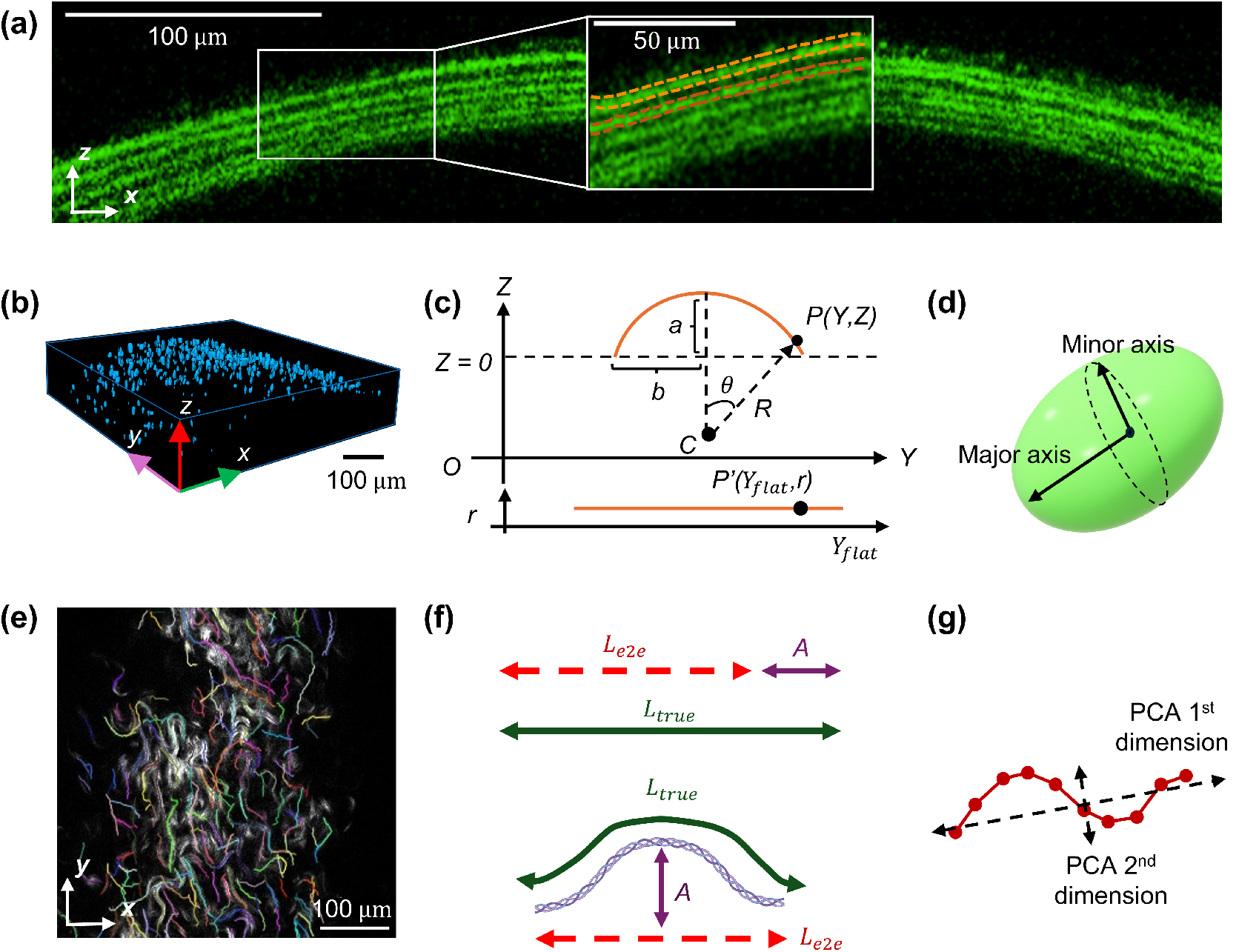
Microstructural analysis workflow. (a) Elastin processing pipeline: elastin channel stacks are reoriented into the *XZ* view and individual lamellae are traced (orange dashed lines) via circle-based geometry (Taubin fitting) to calculate lamellar thickness and interlamellar spacing. (b) 3D segmentation of SYTO-stained nuclei in a representative pressurized and axially extended murine aorta. (c) Schematic illustrating cylindrical flattening of the vessel wall in the *Y Z* plane. *P* (*Y, Z*) is a point on the curved wall and *P* ^*′*^(*Y*_flat_, *r*) is the mapped point in the flattened coordinate system. *C* is the center of the circle approximating wall curvature and *R* is the radius of curvature. *a* is the maximum wall height above *Z* = 0 and *b* is half of the wall span in *Y*, both used to estimate *R. Z*_new_ is the shifted *Z* coordinate used to reference the circular arc. *r* is the radial coordinate of *P* defined as the distance from *C* and preserved under flattening. *θ* is the angular position of *P* about *C. Y*_flat_ is the unwrapped circumferential coordinate. (d) Nuclear shape quantification: segmented nuclei are fit with ellipsoids to determine principal axes, from which major-axis orientation and nuclear aspect ratio (NAR) are derived. (e) Collagen channel stacks are processed using CT-FIRE to extract individual fiber traces and per-slice average fiber widths. (f) Fiber traces are used to calculate amplitude (*A*), end-to-end length (*L*_*e*2*e*_), and true length (*L*_*true*_) of each collagen fiber for quantification of fiber curvature. (g) Fiber orientation is captured by applying principal component analysis (PCA) to individual fiber traces and used to model orientation distributions.

Interlamellar spacing was defined as the lumen-free gap between adjacent lamellar boundaries and calculated from best-fit radii as *g*_*n,n*+1_ = *R*_i, *n*+1_ − *R*_o, *n*_. This metric isolates the separation between lamellae independent of lamellar thickness. All measurements were scaled using a known pixel-to-micron conversion factor.

### 2.4. Nuclear Fluorescence Image Analysis

Orientation and shape of medial vascular smooth muscle cells (VSMCs) and adventitial fibroblasts were derived from 3D multiphoton image stacks of SYTO-stained nuclei. To reduce noise and improve signal quality, image stacks were smoothed using a Gaussian filter [84], followed by background subtraction via morphological opening to correct for uneven illumination. Otsu’s method was then used to determine an optimal threshold for binary conversion, yielding a binarized image stack [85]. Small objects below a minimum volume threshold (*<* 50 pixels) were removed to eliminate residual noise not captured by thresholding. Connected component analysis [86] was then performed to identify individual nuclei and extract their centroid coordinates (Fig. 2b).

To distinguish fibroblasts from VSMCs based on their spatial location within the aortic wall, a cylindrical coordinate transformation was applied to flatten the curved vessel geometry (Fig. 2c). The local radius of curvature was estimated as *R* = (*a*^2^ + *b*^2^)*/*2*a*, where *a* is the maximum range of the image along the *Z* axis and *b* is half the image width in the *Y* direction, both expressed in microns based on axis-appropriate scaling. The axial coordinate was adjusted as *Z*_new_ = *Z* + *R* − *a*. Radial and angular coordinates were then computed as 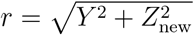 and *θ* = arcsin(*Y/r*). The flattened circumferential coordinate was defined as *Y*_flat_ = *r · θ*. This transformation maps each image-plane point *P* (*Y, Z*) → *P* ^*′*^(*Y*_flat_, *r*), straightening the circumferential direction while preserving the radial coordinate.

In the flattened representation, nuclei were ranked by radial position and classified as VSMCs or fibroblasts based on their location within vessel wall compartments. For each sample, the fractional thickness of the adventitia was determined by visual delineation of collagen-rich regions relative to elastin-rich medial tissue. Centroid coordinates of segmented nuclei were computed in physical units (*μ*m) as 3D-connected-component centroids using the built-in function *regionprops3*. Nuclear class labels (VSMCs vs. fibroblasts) and centroid coordinates were then mapped back to the original 3D space for morphological analysis.

Each nucleus was isolated as a 3D object, and a surface mesh was generated using a marching cubes algorithm (isovalue 0.5). Vertex coordinates were converted to physical units (*μ*m) and mean-centered. Principal axes were then obtained as the eigenvectors of the vertex covariance matrix, with square roots of the eigenvalues proportional to the semi-axis lengths. This ellipsoidal fit defined the major and minor axes of each nucleus, as illustrated in Fig. 2d.

Nuclear orientation was quantified using the major-axis eigenvector, which was projected into the imaging plane and converted to an undirected in-plane angle *θ* ∈ [0, *π*), such that *θ* and *θ* + *π* represent the same axis. Orientation distributions were then obtained by binning *θ* values separately for VSMCs and fibroblasts and normalizing the resulting histograms to unit area.

Nuclear aspect ratio was defined as *NAR* = Major Axis Length*/*Minor Axis Length and served as a scale-invariant measure of nuclear elongation. NAR values were computed for all nuclei, and descriptive statistics were calculated separately for VSMCs and fibroblasts.

### 2.5. Collagen Second Harmonic Generation Image Analysis

Collagen architecture was characterized from second harmonic generation (SHG) images acquired in *XY* optical planes spanning the aortic wall. In the descending thoracic aorta, collagen fibers are predominantly oriented within the imaging plane (anatomical circumferential and axial directions), permitting reliable in-plane orientation analysis from individual optical sections [57, 56, 68]. Collagen fiber traces were extracted using CT-FIRE [62, 63], which enhances elongated features via a curvelet-based transform followed by edge detection (Fig. 2e). Default CT-FIRE settings were applied uniformly across samples. Traces shorter than 50 pixels (23.5*μ*m) were excluded to minimize noise-driven false positives and eliminate incompletely captured fibers.

#### 2.5.1. Fiber Straightness and Amplitude

Adapting published methods [9, 69, 70], fiber straightness was defined as *P*_*s*_ = *L*_*e*2*e*_*/L*_true_ and represents the ratio of the end-to-end distance *L*_*e*2*e*_ to the traced fiber length *L*_true_. A value of *P*_*s*_ = 1 corresponds to a perfectly straight fiber, while values approaching 0 indicate greater waviness. Within each sample, straightness distributions were fitted to a Beta probability density function,

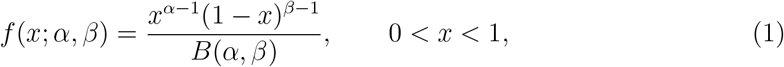

where *x* ≡ *P*_*s*_, *α* and *β* are shape parameters, and

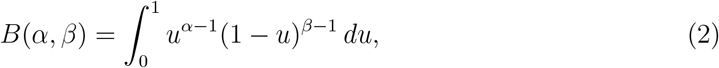

is the normalization constant, where *u* serves as a dummy integration variable. Parameters *α* and *β* were estimated using the built-in betafit function [9, 69, 70]. When holding *β* fixed, larger *α* shifts probability mass toward *x* ≈ 1, indicating higher prevalence of straighter fibers. Conversely, larger *β* at constant *α* shifts probability mass toward *x* ≈ 0, indicating greater waviness. When *α >* 1 and *β >* 1, the distribution is unimodal on (0, 1), while *α <* 1 and *β <* 1 yield a U-shaped distribution with increased probability near the bounds. The mean straightness satisfies 𝔼[*x*] = *α/*(*α* + *β*), such that increases in *α* or decreases in *β* raise the average straightness.

Fiber amplitude was defined as *A* = *L*_true_ − *L*_*e*2*e*_ and represents the excess traced length relative to end-to-end distance (Fig. 2f) [69]. Fibers were further classified as circumferential or axial based on their orientation relative to the dominant direction (threshold *±*40^°^), and straightness and amplitude metrics were computed separately for each group.

#### 2.5.2. Network Porosity and Linear Density

Collagen network porosity was analyzed using 3D reconstructions derived from CT-FIRE traces. For each *Z*-plane, 2D fiber traces were converted into binary images and dilated to the mean fiber width reported by CT-FIRE per optical section. Binary images were then stacked to form a volumetric representation of the 3D network.

Porosity was calculated as *𝒫* = *V*_empty_*/V*_total_ and represents the fraction of non-fiber voxels *V*_empty_ relative to the total voxel count *V*_total_. As a complementary metric, linear density was calculated as *ρ*_*L*_ = *L*_total_*/V*_total_, where *L*_total_ is the total traced fiber length (pixels).

#### 2.5.3. Fiber Orientation

A single in-plane orientation angle was computed for each traced collagen fiber using principal component analysis (PCA) of the *XY* centerline coordinates, with the primary eigenvector defining the dominant fiber axis (Fig. 2g). Orientation angles *θ* were then pooled across all fibers within a specimen and binned with a resolution of Δ*θ* = 0.5^°^ over the range (0°–180°), or (0–*π*) in radians.

Let *n*(*θ*_*i*_) denote the number of fibers in the *i*-th angular bin centered at *θ*_*i*_ and let *N* = Σ_*i*_ *n*(*θ*_*i*_) be the total number of fibers. The empirical probability density of fiber orientations was defined as

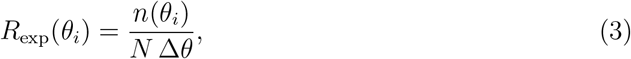

such that

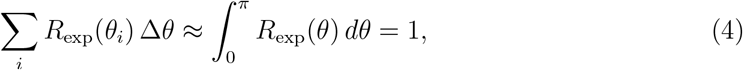

where *R*_exp_(*θ*) represents the probability density of observing a fiber with in-plane orientation angle *θ*.

#### 2.5.4. Von Mises Model Fitting

Empirical distributions of collagen fiber orientations were consistent with an isotropic fiber network superimposed with anisotropic fiber families. Following prior work [65, 66, 67], measured orientation densities were modeled by a two-fiber-family von Mises mixture,

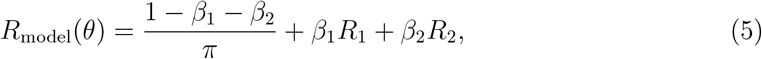

where (1−*β*_1_ −*β*_2_)*/π* represents the isotropic background of randomly oriented fibers [67], while the coefficients *β*_1_ and *β*_2_ weight the respective contributions of the two fiber families *R*_1_ and *R*_2_, such that 0 ≤ *β*_1_ + *β*_2_ ≤ 1. Each fiber family *R*_*i*_ (*i* = 1, 2) is described by

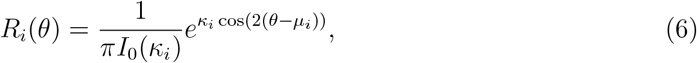

where *I*_0_ is the modified Bessel function of the first kind, order zero, *μ*_*i*_ is the preferred orientation, and *κ*_*i*_ is the concentration parameter controlling peak sharpness, with larger *κ*_*i*_ corresponding to tighter alignment around *μ*_*i*_. The use of cos 2(*θ* − *μ*_*i*_) enforces *π*-periodicity of *R*_*i*_(*θ*), consistent with head-tail symmetry in fiber orientations, where *θ* and *θ* + *π* describe the same fiber axis.

Because the weights *β*_*i*_ govern the relative contributions of the anisotropic components, the ratio *β*_1_*/β*_2_ was used to quantify asymmetry between fiber families. When *κ*_*i*_ are small or *μ*_1_ and *μ*_2_ are closely spaced, the two fiber families broaden and overlap, producing an approximately flat composite distribution. Explicit inclusion of the isotropic term thus allows discrimination between this limiting scenario and a truly uniform background. In addition, the isotropic term ensures that *R*_model_(*θ*) integrates to unity over [0, *π*).

Model parameters were estimated by minimizing a normalized mean square error,

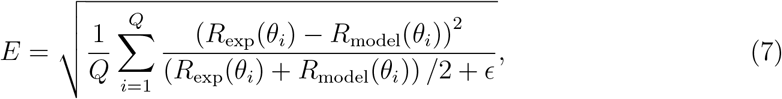

where *Q* is the number of angular bins, *θ*_*i*_ denotes the center of the *i*-th bin, and *ϵ* is a small regularization constant introduced to prevent numerical instability. Optimization was performed subject to constraints 0 ≤ *β*_1_, *β*_2_, 0 ≤ *β*_1_ + *β*_2_ ≤ 1, and 0 ≤ *μ*_1_, *μ*_2_ ≤ *π*. Best-fit distributions were renormalized to unit area.

#### 2.5.5. Volume Fraction Analysis

Collagen and elastin volume fractions were quantified from multiphoton image stacks using a framework adapted from a prior developmental study in murine aortic tissues [68]. The process encompasses two stages: first, the derivation of fixed, robust, per-channel thresholds from a balanced training subset; second, the batch computation of volume fractions across all samples using those thresholds.

In the first stage, a subset of samples (*n* = 3) was selected. This training subset was considered balanced in that it spanned the range of imaging conditions present in the entire dataset, while avoiding over-representation of any single acquisition. Specifically, the three samples were chosen to provide comparable vessel wall coverage and signal quality across image stacks and to include representative intensity distributions in both channels. Selection was performed after quality control screening to exclude volumes with incomplete wall coverage, gross saturation or clipping after normalization, missing slices, or severe motion artifacts.

Each dataset contained co-registered 3D TIFF volumes corresponding to collagen (SHG) and elastin (autofluorescence) signals. For each sample and channel, the 3D volumes were cropped or permuted to a common *Z × Y × X* size. Voxel intensities were then linearly rescaled on a per-volume, per-channel basis. Let *I*(*x, y, z*) denote the raw intensity at voxel (*x, y, z*). For any given channel, *I*_min_ and *I*_max_ were defined as the minimum and maximum voxel intensities within the cropped 3D volume. The normalized intensity was then computed as

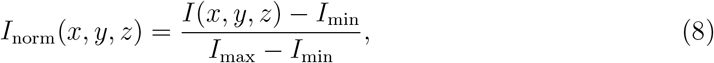

which placed *I*_norm_ in the range [0, 1]. Values numerically outside this range due to interpolation or rounding were clipped to [0, 1].

Binary masks for each channel were obtained using Otsu’s method [85] and then combined to form a vessel wall mask via morphological closing and hole filling. Per-image Otsu thresholds for collagen (*t*_*collagen*_) and elastin (*t*_*elastin*_), alongside an additional 75th-percentile cutoff of nonzero voxels, were applied to yield median thresholds across the training set.

In the second stage, these median thresholds were imposed uniformly across all images to generate binary collagen and elastin segmentations. The union of these masks defined the vessel wall region of interest, followed by morphological closing with a spherical structuring element (radius 1 voxel) and 3D hole filling to obtain a contiguous mask. Voxels present in both channels were attributed to the constituent with the highest normalized intensity. Volumes of collagen (*V*_collagen_) and elastin (*V*_elastin_) were computed as the number of voxel assigned to each constituent within the vessel wall mask, multiplied by the physical voxel volume. Voxels not labeled as collagen or elastin were grouped into a residual volume component, reflecting voxels introduced by morphological operations on the union mask and low-intensity wall regions below both channel thresholds. Total vessel wall volume (*V*_vessel_) was calculated in a similar fashion from the final wall mask. Volume fractions for collagen, elastin, and unsegmented wall constituents were defined as *φ*_collagen_ = *V*_collagen_*/V*_vessel_, *φ*_elastin_ = *V*_elastin_*/V*_vessel_, and *φ*_other_ = 1 − *φ*_collagen_ − *φ*_elastin_, respectively. These fractions quantify the relative contributions of different microstructural constituents to the vessel wall volume.

## 3. Results

### 3.1. Elastin

In the reoriented *XZ* views, elastic lamellae were sharply resolved across all specimens, providing a reliable basis for quantifying their geometric features. Circle-based fitting produced concentric inner and outer arcs with midline segments for each lamella. Because this approach is curvature-aware in the *XZ* plane, the resulting measurements are less sensitive to field-of-view tilt or local wall curvature, compared to radial distances extracted from raw *XY* stacks. Using these fits, we observed modest layer-to-layer variability in both elastic lamellar thickness and interlamellar spacing, consistent with prior reports describing the organization of medial elastic lamellar units in thoracic aortic tissues [15, 55, 23]. Values of both parameters for the first five elastic lamellae from a representative aorta under physiological luminal pressurization and axial extension are reported in Table 1.

**Table 1.** Geometrical features of elastic lamellae in a representative pressurized and axially extended murine aorta with wall thickness of 35.68 *μm*. Lamellae are numbered sequentially from the luminal side of the tunica media, with 1 corresponding to the internal elastic lamina.

| Lamella Number | Lamellar Thickness ($\mu m$) | Interlamellar Distance ($\mu m$) |
| --- | --- | --- |
| 1 | 6.28 | 2.65 |
| 2 | 3.49 | 3.62 |
| 3 | 4.95 | 2.95 |
| 4 | 3.93 | 3.58 |
| 5 | 4.24 | — |

### 3.2. Nuclear Deformation

Implementation of the cylindrical flattening and partitioning workflow revealed distinct distributions of nuclear major-axis orientation between medial and adventitial cell populations under physiological luminal pressurization and axial extension (Fig. 3a for a representative aorta). Medial nuclei exhibited pronounced circumferential alignment, while adventitial nuclei showed a broader orientation distribution. This pattern reflects layer-specific cellular organization within the aortic wall. Through mechanical coupling between the actin cytoskeleton and the nucleus, cytoskeletal forces can deform the nuclear envelope, such that isotropic loading primarily alters nuclear volume, whereas directionally biased tension produces sustained nuclear elongation [8, 87]. Spindle-shaped VSMCs aligned with medial lamellar units generate and dynamically regulate cytoskeletal tension primarily along the circumferential axis, through basal contractile tone and responses to cyclic deformation of the surrounding ECM. In contrast, adventitial fibroblasts, characterized by more heterogeneous morphologies and orientations, lack a consistent dominant axis for force transmission.

**Figure 3.**
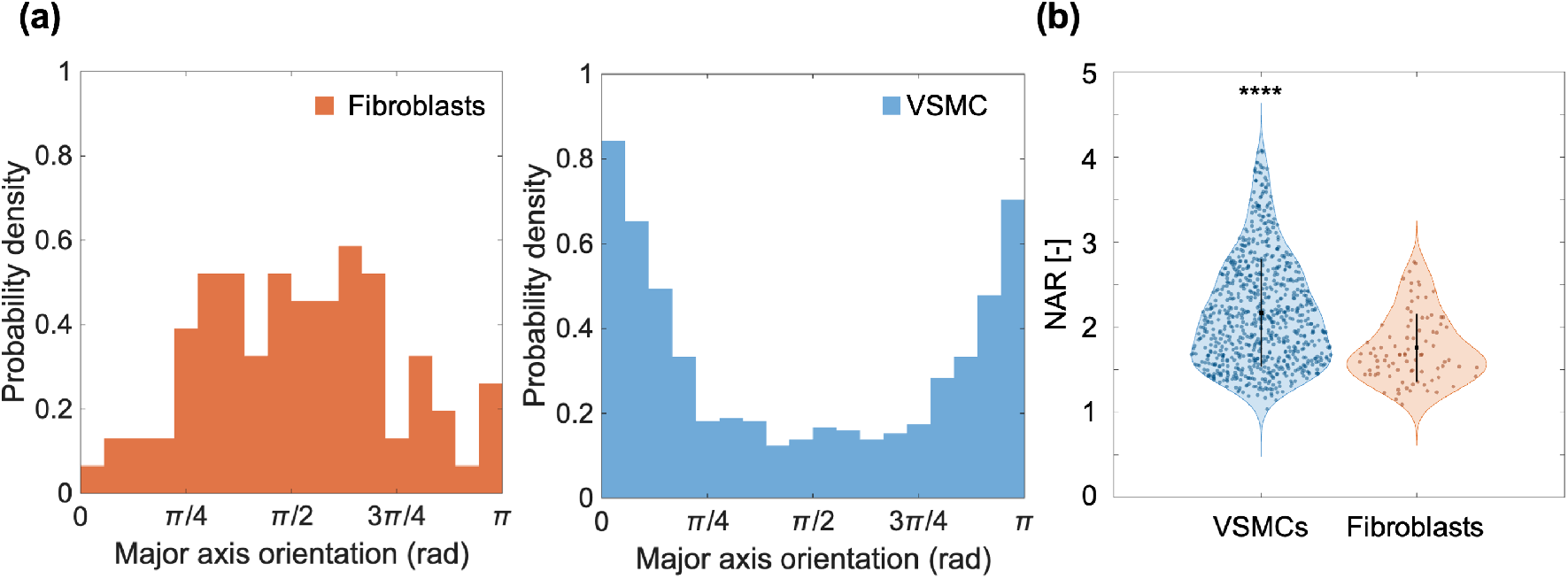
(a). Normalized probability density distributions of nuclear major-axis orientation for VSMCs and fibroblasts over [0, *π*], with 0 and *π* corresponding to the circumferential direction, and *π/*2 corresponding to the axial direction. Data from a representative pressurized and axially extended murine aorta. (b). Distribution of NAR for VSMCs in the media and fibroblasts in the adventitia for the same representative sample. Violin plots illustrate the kernel density of NAR values. The filled black dot denotes the mean (*μ*) and vertical bars extend to one standard deviation (SD) above and below *μ* in each plot. *μ* = 2.17 and SD = 0.634 for *n* = 789 VSMCs. *μ* = 1.75 and SD = 0.401 for *n* = 88 fibroblasts. Between group difference was assessed using a one sided Wilcoxon rank sum test (Mann–Whitney U) with the alternative hypothesis that VSMC NAR exceeded fibroblast NAR (*p* = 7.2 *×* 10^−10^), as indicated by the asterisks.

Under these orientation-dependent loading conditions imposed by cell-type–specific morphology and cytoskeletal alignment, VSMCs exhibited higher NAR values than fibroblasts (Fig. 3b for the same representative sample).

### 3.3. Collagen Network

#### 3.3.1. Fiber Straightness and Amplitude

CT-FIRE traces provided per-fiber centerlines, from which straightness *P*_*s*_ and amplitude *A* were computed. Empirical distributions of *P*_*s*_ were well described by Beta(*α, β*) fits, yielding compact sample-level descriptors within the bounded interval [0, 1] (Fig. 4a for a representative aorta). As expected, these distributions were skewed toward higher *P*_*s*_ values, reflecting fiber recruitment and straightening under under physiological luminal pressurization and axial extension, with a low-probability tail at smaller *P*_*s*_ corresponding to wavier fiber segments. Because both *P*_*s*_ and *A* depend on trace truncation at fiber crossings and the minimum-length filter, we used default CT-FIRE parameters and applied a conservative 50-pixel cutoff to limit short, spurious segments. These modeling choices prioritize comparability across samples over maximal trace yield.

**Figure 4.**
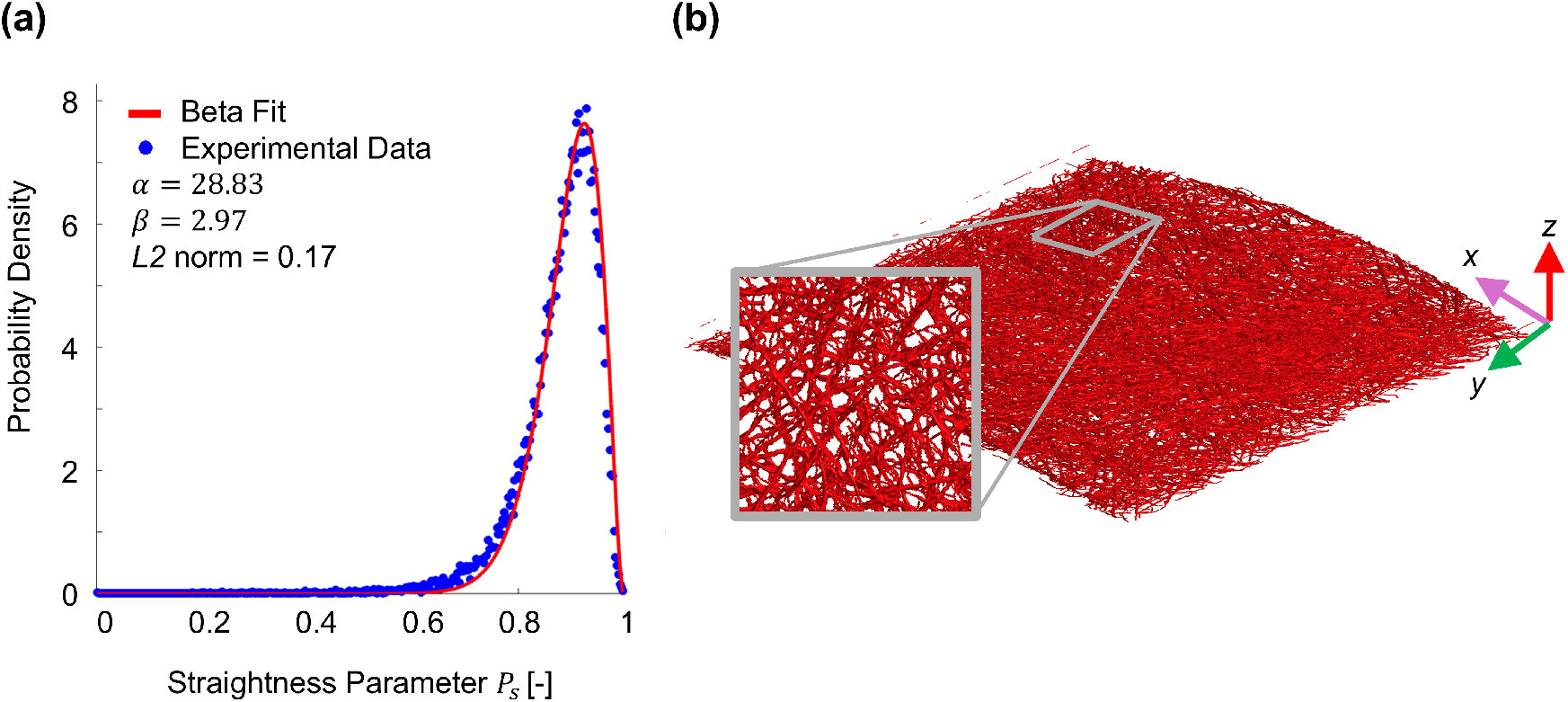
(a). Beta–distribution fit of fiber straightness parameter *P*_*s*_ in a representative pressurized and axially extended murine aorta. Blue symbols are experimental data points and the red curve is the best-fit Beta(*α, β*) probability density function. Text annotations report Beta distribution parameters and *L*2 norm error for the fitting. (b). 3D reconstruction of the collagen fiber network for a representative pressurized and axially extended murine aorta. The gray rectangle marks a region of interest, while the inset shows a magnified view of the same area to highlight local branching and bundling. Axes indicate the imaging coordinate frame. Porosity (void fraction) and linear density (total centerline length per unit volume) were computed from this reconstruction.

#### 3.3.2. Porosity and Linear Density

A pragmatic 3D approximation of the collagen network was obtained by stacking per-plane fiber traces, dilated to the mean fiber width at each *Z* location. The resulting binary volume enabled direct computation of porosity (void fraction) and linear density (total trace length per unit volume). In a representative reconstruction under physiological loading conditions (Fig. 4b), the collagen network occupied a minority of voxels, yielding high porosity (*𝒫* = 0.89), yet retained substantial centerline length per unit volume (*ρ*_*L*_ = 0.06*μ*m^−2^), indicative of dense in-plane packing with limited through-thickness branching. This architecture is consistent with prior reports describing predominantly in-plane collagen fibers in the thoracic aortic media [57, 56, 68, 55, 15,,23, 9]. Because porosity depends on the imposed dilation (mean width), we recommend that within-study comparisons be performed using identical reconstruction parameters.

#### 3.3.3. Fiber Orientation

Two-family von Mises mixtures models resolved dominant orientation modes across all specimens, while also accounting for dispersed/isotropic content through the constant term. In representative fits (Fig. 5a,b, for two pressurized and axially extended aortic samples), the model captured both peak locations (mean directions *μ*_*i*_) and dispersion (concentrations *κ*_*i*_), while maintaining normalized orientation densities across the interval [0, *π*].

**Figure 5.**
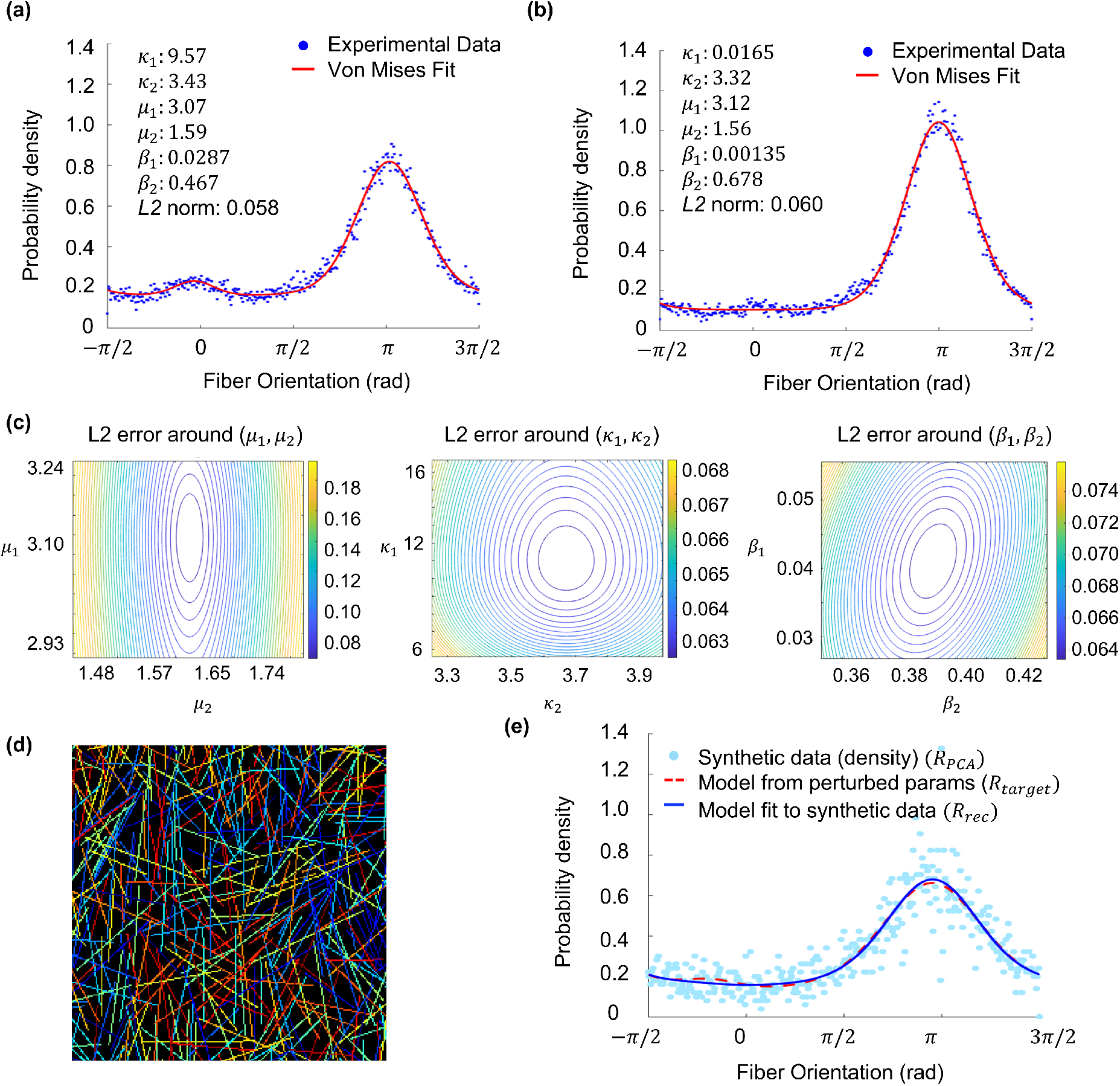
Collagen fiber orientation analysis. (a–b) Representative von Mises mixture model fits for two pressurized and axially extended murine aortic samples imaged under identical conditions. Blue circles show the fiber orientation density measured experimentally, while red lines represent the best fit obtained from a von Mises mixture model. Each panel includes text annotations listing best-fit parameters (concentrations *κ*_*i*_, mean directions *μ*_*i*_ in radians, weights *β*_*i*_) and the *L*2 error norm. Despite inter-specimen differences in peak location and sharpness, including relatively broad or low-amplitude peaks, the model well captures the dominant orientations and degree of alignment. (c) Local *L*2 error landscapes for either (*μ*_1_, *μ*_2_), (*κ*_1_, *κ*_2_), or (*β*_1_, *β*_2_), with all other parameters held constant. Convex contours centered near the optimum indicate a locally unique solution. (d) Binary synthetic image generated by sampling orientations from the best-fit von Mises mixture model in (a). Colors are used solely to distinguish individual fibers and carry no quantitative meaning, while the underlying image is binary, as in the experimental data. The synthetic field recapitulates dominant orientations, dispersion about the modes, and crossing density observed experimentally. (e) Forward-inverse validation of the two-family von Mises orientation model. Light blue circles show the PCA-derived orientation density from the synthetic data (*R*_PCA_), the red dashed line denotes the target model used to generate the synthetic field (*R*_target_), and the blue line indicates the re-fitted model obtained from the synthetic data (*R*_rec_).

To visually assess local robustness and parameter identifiability, we mapped the objective landscape in the neighborhood of the optimum by computing error contours for parameter pairs, while holding the remaining four parameters at their best-fit values. The error metric was defined as the continuous *L*2 norm between the normalized experimental density and the model density,

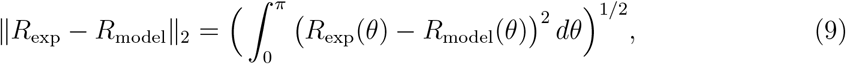

such that each contour map visualized how the misfit changed with joint perturbation of two parameters.

We examined three parameter pairs that captured the main couplings in the model: (i) the fiber family weights (*β*_1_, *β*_2_), subject to 0 ≤ *β*_1_, *β*_2_ and *β*_1_ + *β*_2_ ≤ 1; (ii) the mean fiber family directions (*μ*_1_, *μ*_2_) with *π*–periodicity enforced via cos(2(*θ* − *μ*_1_)) and cos(2(*θ* − *μ*_2_)); and (iii) the fiber family concentration parameters (*κ*_1_, *κ*_2_), with *κ*_1_, *κ*_2_ *>* 0. For (*β*_1_, *β*_2_) and (*κ*_1_, *κ*_2_) we used multiplicative neighborhoods of *±*10% around the optimum, whereas for (*μ*_1_, *μ*_2_) we used additive windows of *±*10^°^. For each pair of parameter values, the model was re-normalized and the *L*2 error was re-computed.

The resulting contour maps were locally convex for all samples, with the optimum located at or near the lowest error level, consistent with a unique and stable minimum within these neighborhoods (Fig. 5c for a representative pressurized and axially extended aorta). The shape of the contour levels further reflected the sensitivity of the model fit to changes in the parameter values and the degree of coupling between parameters. Nearly circular contours indicated comparable sensitivity of the fit to both parameters, such that variations in one parameter could not be compensated by adjusting the other without degrading the fit. In contrast, elliptical contours elongated along one axis indicated lower sensitivity of the fit in that direction, mirroring limited compensatory behavior in which changes on one parameter could be partially offset by adjustments in the other, while maintaining a similar fit.

To complement the visual contour analysis, we evaluated the local Hessian of the *L*2 error surface at the optimum for each parameter pair. The Hessian quantifies the local curvature of the misfit with respect to parameter perturbations, with eigenvalues describing curvature along the principal directions in the parameter space. Consistent with visual inspection of the contour maps, all samples exhibited positive eigenvalues, confirming that the optimum corresponded to a locally convex minimum. We additionally computed the Hessian condition number, defined as the ratio of the largest to smallest eigenvalue, as a measure of anisotropy in local curvature. In agreement with the contour shapes observed visually, modest condition numbers indicated comparable curvature and similar identifiability of the paired parameter, while larger values revealed directions of reduced constraint within parameter pairs. For the representative specimen shown in Fig. 5c, Hessian eigenvalue ratios (largest/smallest) were 2.8 for (*β*_1_, *β*_2_), 50.5 for (*μ*_1_, *μ*_2_), and 1.0 *×* 10^3^ for (*κ*_1_, *κ*_2_), computed from the corresponding eigenvalues pairs [25.6, 9.16], [22.5, 0.444], and [6.8 *×* 10^−2^, 6.8 *×* 10^−5^].

To evaluate the image relevance of the von Mises parameters – that is, how well they encoded the fiber organization observed in the image set used for their identification – we implemented a combined forward–inverse validation using synthetic microscopy. This approach mirrored the real-data pipeline, including manual peak seeding, to ensure that any improvements or biases arose from the data rather than from changes in the workflow. The validation was performed on a von Mises mixture model best fit to the distribution of collagen fiber orientations in a representative aortic sample under physiological loading.

First, we tested the forward mapping from parameters to image. Starting from the optimal parameter set, we generated a target fiber orientation density distribution *R*_target_(*θ*) by applying small multiplicative perturbations to *κ*_*i*_ and *β*_*i*_, along with additive *±* jitter to *μ*_*i*_. We then sampled a large set of fiber orientations *θ*_*j*_ from *R*_target_. Next, we constructed a binary synthetic image by rendering idealized fiber elements as slender rectangles of fixed thickness with random length and position, aligned with the sampled orientations (Fig. 5d). These synthetic fibers were written into a label matrix with one unique label per fiber, yielding both a composite binary image and per-fiber masks.

Second, we evaluated the inverse mapping from image back to parameters. For each labeled synthetic fiber, we extracted the major axis direction via PCA of the pixel coordinates (principal eigenvector) and converted it to an in plane principal axis orientation angle in the range [0, *π*), measured relative to the image *X* axis. Binning these angles on the same grid as the model yielded a normalized, PCA-derived fiber orientation density distribution *R*_PCA_(*θ*). We then repeated the fitting workflow applied to the real data, including *R*_PCA_ smoothing, manual selection of two peaks to seed *μ*_1_ and *μ*_2_, enforcement of the constraints 0 ≤ *β*_1_, *β*_2_, *β*_1_ + *β*_2_ ≤ 1, and *κ*_1_, *κ*_2_ *>* 0, and parameter re-fitting through minimization of the continuous *L*2 misfit. This produced a recovered model for the fiber orientation density distribution *R*_rec_(*θ*) (Fig. 5e).

Agreement between recovered and target values was quantified as

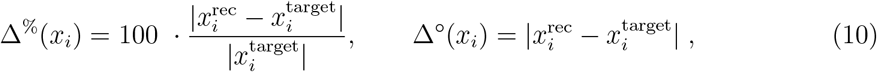

where *x*_*i*_ denotes a model parameter, Δ^%^ quantifies relative percentage differences for scalar parameters, and Δ^°^ measures angular deviations of the mean fiber directions, expressed in radians. For the validation example considered here, parameter-level deviations were Δ^%^(*β*_1_) = 69%, Δ^%^(*β*_2_) = 11%, Δ^%^(*κ*_1_) = 314%, Δ^%^(*κ*_2_) = 21%, Δ^°^(*μ*_1_) = 0.05 rad, and Δ^°^(*μ*_2_) = 0.01 rad. Although some discrepancies are larger than one may expect, these values should be interpreted with care. Differences between recovered parameters and their targets are informative in that they reveal which aspects of the model are tightly constrained by the data and which are susceptible to ambiguity or compensatory trade-offs. In this context, large deviations do not necessarily imply poor recovery, but may instead reflect inherent identifiability limits or structured parameter interactions within the model. We therefore interpret the parameter-level results through this lens before discussing the implications of individual deviations.

Discrepancies in *μ*_*i*_ provide a direct measure of directional fidelity. In this validation instance, the recovered mean directions showed strong agreement with their targets, reflecting robust recovery of fiber orientation. By contrast, large angular discrepancies accompanied by small percentage differences in *κ*_*i*_ and *β*_*i*_ would point to uncertainty in peak location.

Percentage deviations in *β*_*i*_ and *κ*_*i*_ primarily reflect parameter identifiability within the von Mises mixture model rather than inaccuracies in the recovered orientation distribution, particularly when a component weight *β*_*i*_ is small. In this validation example, the largest deviations were associated with a low-weight fiber family, such that variations in these parameters exerted minimal influence on the overall fit. Although *κ*_*i*_ governs peak sharpness, its contribution to the mixture is scaled by the corresponding weights *β*_*i*_. As a result, when *β*_*i*_ is small, relatively large percentage deviations in either parameter can occur without materially altering the distribution. By contrast, comparable deviations in highly weighted fiber families would point to ambiguity in peak amplitude or dispersion and warrant closer scrutiny. Similarly, large Δ^%^(*β*_*i*_) values occurring along shallow error surfaces in the (*β*_1_, *β*_2_) plane would reflect compensatory redistribution of weight between fiber families with minimal impact on fit quality.

Distribution-level errors should therefore be interpreted in conjunction with parameter-level deviations. Visual inspection of the fiber orientation density distributions in this validation example revealed close alignment of *R*_PCA_ with *R*_target_ and of *R*_rec_ with *R*_PCA_. This qualitative agreement was supported by negligible distribution-level errors, with *L*2(*R*_target_, *R*_PCA_) = 0.153 and *L*2(*R*_rec_, *R*_PCA_) = 0.152, suggesting that most of the observed error arises from the forward validation step. Near-equal *L*2 values despite noted differences in *β*_*i*_ and *κ*_*i*_ are attributable to regions of the parameter space with limited sensitivity and intrinsic weight–concentration trade-offs across fiber families, rather than substantive differences in the recovered orientation distribution. By contrast, large angular deviations occurring at stable *L*2 would more typically signal ambiguity in peak location rather than a true structural mismatch.

Taken together, findings from this test case validation confirm that von Mises parameters inferred from real images encode measurable image-level structure that can be rendered, remeasured, and reliably re-identified.

#### 3.3.4. Volume Fraction

Volume fractions for collagen, elastin, and the remaining vessel wall content were computed using fixed, channel-specific intensity thresholds applied uniformly across samples. In a representative murine aortic specimen under luminal pressurization and axial extension, voxel-wise measurements from co-registered 3D multiphoton stacks yielded *φ*_collagen_ = 0.25, *φ*_elastin_ = 0.47, and *φ*_other_ = 0.28 relative to the total vessel wall volume. These values are consistent with compositional analyses of two-dimensional histological cross-sections from fixed murine aortic tissues. For reference, the descending thoracic aorta of 20-week-old male wildtype mice on a mixed genetic background was reported to comprise approximately 32% elastin and 39% collagen by area, with the remaining fraction attributed to non-fibrillar constituents such as smooth muscle and proteoglycan-rich matrix [88]. Using Otsu-based thresholds and voxel/pixel-count definitions, a complementary multiphoton-based approach further quantified layer-specific volume fractions in aortic tissues from female wildtype mice throughout development, yet direct comparison with our results is precluded by differences in the treatment of total vessel wall volume [68].

Because *φ* is computed by counting all voxels within the segmented vessel wall, we systematically evaluated the sensitivity of these measurements to segmentation and processing parameters. Robustness checks for volume fraction estimates were performed using 3D synthetic datasets for which the ground-truth (GT) collagen and elastin labels were known (Fig. 6a for a representative rendering). Each phantom comprised paired collagen and elastin volumes with GT labels and was processed through a forward imaging model designed to mimic experimental artifacts, including depth-dependent attenuation, linear channel mixing (cross-talk), Poisson and Gaussian noise, salt-and-pepper outliers, blur, shallow intensity gradients along the *Z* axis, striping, and sparse hot pixels (Fig. 6b). Volume fraction measurements for the synthetic data were then obtained by applying the same fixed thresholding, mask-formation steps, and overlap-resolution rules used for the experimental data.

**Figure 6.**
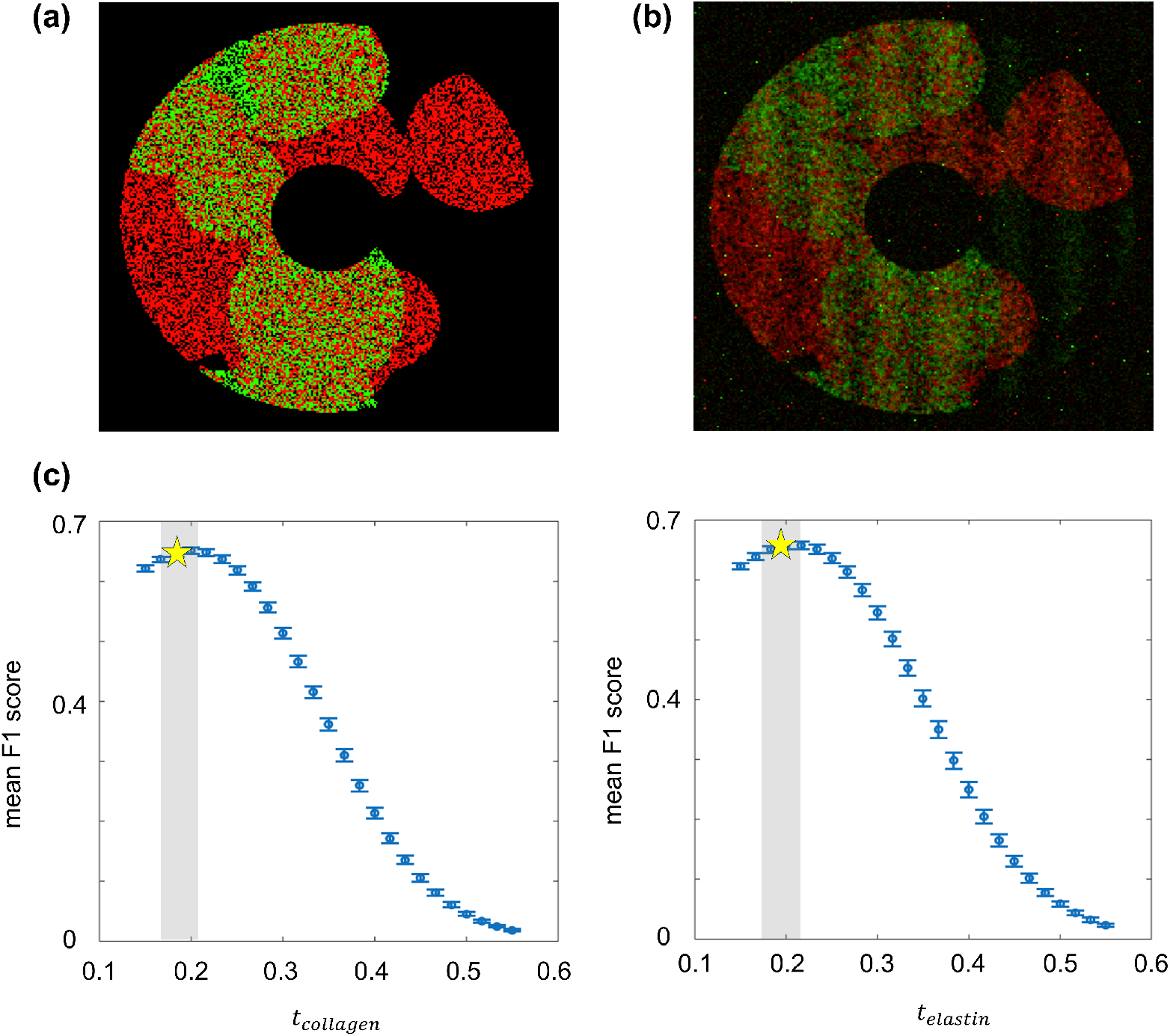
Evaluation of the volume-fraction pipeline using synthetic data. (a) Max *Z* projections of ground-truth label map from a representative synthetic phantom showing collagen (red), elastin (green), and background (black). (b) Corresponding simulated image after including depth attenuation, spectral cross-talk, blur, noise, and intensity gradients. (c) Threshold sweep curves showing mean F1 score versus fixed threshold for collagen (*t*_*collagen*_) and elastin (*t*_*elastin*_), computed on synthetic test volumes with known ground truth labels. Symbols and error bars denote the mean and standard deviation of F1 scores across volumes. The yellow star marks the fixed per-channel threshold obtained from median Otsu values, indicating that the operating point lies within a stable, high-performance threshold region.

First, we quantified the sensitivity of *φ* to the selection of the sample subset from which the fixed thresholds were derived. Using the synthetic data, we performed five-fold crossvalidation, in which thresholds were re-estimated from four-fifths of the phantoms and then applied to the held-out phantom in each fold. We defined the volume fraction drift as *φ*_drift_ = |*φ*_fold_ − *φ*_baseline_|, where *φ*_baseline_ is obtained using thresholds derived from the full phantom set, and *φ*_fold_ is the volume fraction computed for the held-out phantom, using thresholds estimated from the corresponding four-fifths training subset in that cross validation fold. Across folds, the median *φ*_drift_ was ≤ 0.003 for both collagen and elastin, indicating that volume fraction measurements were essentially insensitive to the choice of training subset. We further evaluated segmentation robustness on the same synthetic phantoms via threshold-sweep analyses using the F1 score, defined as the harmonic mean of precision and recall. [89]. Because it penalizes both false positive and false negative voxel assignments, the F1 score is well suited for class-imbalanced voxel classifications such as vessel wall segmentation. F1 values remained stable over a broad range around the chosen thresholds (Fig. 6c) and peaked near the median Otsu threshold selected for collagen (*t*_collagen_ = 0.184) and elastin (*t*_elastin_ = 0.192). The extended peak in F1 indicates that moderate deviations in *t*_collagen_ and *t*_elastin_ produce minimal changes in segmentation quality, supporting the use of fixed study-wide thresholds for estimating volume fractions that are robust to minor threshold variations.

We next examined the influence of vessel wall mask morphology, which was constructed as the union of the thresholded collagen and elastin masks, followed by 3D morphological closing and hole filling. In the synthetic phantoms, doubling the closing radius from 1 to 2 voxels produced negligible changes in either *φ*_collagen_ or *φ*_elastin_, demonstrating that plausible variations in mask-cleanup parameters do not significantly affect volume fraction analysis.

Finally, we assessed the impact of spectral cross-talk arising from imperfect emission filtering, which leads to partial overlap between raw collagen and elastin signals [52, 51, 57].

We modeled this phenomenon as a linear mixing process

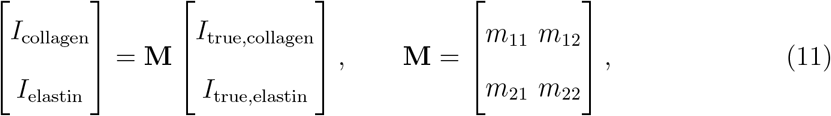

where **M** is a 2 *×* 2 mixing matrix relating measured raw signal intensities *I* to true unmixed intensities *I*_true_ for collagen and elastin [58]. Spectral unmixing was performed by applying **M**^−1^ to recover separated channels.

Because **M** is estimated from calibration images and may contain small errors, we tested the sensitivity to calibration uncertainties by perturbing each entry of **M** by *±*10% in the synthetic phantoms, and recomputing volume fractions following unmixing, while holding all thresholds fixed. We defined the mixing sensitivity as Δ*φ*_*cal*_ = |*φ*(**M**_Δ_) − *φ*(**M**)|, where *φ*(**M**) denotes the volume fraction obtained using the nominal calibration matrix and *φ*(**M**_Δ_) denotes the estimate after unmixing with a perturbed matrix **M**_Δ_. Sensitivity measures were averaged for all synthetic phantoms and independent perturbations applied to the entries of **M** to yield mean and dispersion parameters across realizations. Under these conditions, the mean Δ*φ*_*cal*_ was *<* 0.040 for collagen and *<* 0.057 for elastin, indicating that moderate calibration-level errors in **M** produce only small absolute changes in volume fraction estimates.

To quantify the net impact of cross-talk on volume fraction accuracy relative to GT, we computed the absolute difference in volume fractions referenced to GT as 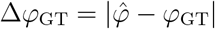, where 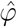 was obtained by applying the full analysis pipeline to mixed, noisy synthetic imaging data and *φ*_GT_ was computed directly from the phantom labelled volumes. Even under conditions in which spectral cross-talk degraded per-voxel segmentation quality (F1 ≈ 0.67), GT-referenced differences remained modest, with Δ*φ*_GT_ *<* 0.045 for collagen and Δ*φ*_GT_ *<* 0.050 for elastin.

Together, these analyses show that the fixed-threshold volume fraction pipeline yields stable and reproducible estimates, and that combining conservative, study-wide thresholding with basic channel separation enhances the robustness of quantitative multiphoton image analysis [51, 68, 63].

## 4. Concluding Remarks

In summary, we developed an end-to-end workflow to quantify aortic wall microstructure from co-registered SHG and TPEF image stacks. This pipeline supports constituent-specific quantification of ECM architecture and nuclear morphology by integrating curvature-aware measurements with segmentation following cylindrical wall flattening.

Application of the workflow yielded several key outcomes. First, elastic lamellar thickness and interlamellar spacing were quantified reliably from *XZ* views despite vessel wall curvature. Second, curvature correction allowed for differences in nuclear shape and orientation to emerge between resident cell populations in the media and adventitia. Third, a Beta distribution effectively captured collagen fiber straightness measured with CT-FIRE. Fourth, a von Mises mixture model incorporating an isotropic term and two fiber families succinctly described collagen fiber orientation with stable parameters. Fifth, dilation-based 3D reconstructions supported estimation of porosity and linear fiber density assuming inplane collagen fiber architecture. Finally, fixed-threshold volume fraction estimates were robust to modest variation in threshold selection, vessel masking, and spectral cross-talk.

A defining strength of the workflow is the emphasis on reproducibility, achieved through fixed decision rules, standardized preprocessing steps, and thorough validation on synthetic datasets. This design enables application to different samples without manual adjustment or case-by-case tuning, thereby supporting meaningful comparisons across biological conditions. Despite these advantages, the current implementation carries some limitations, as assignment of nuclei to medial and adventitial compartments is based on predefined rules, and collagen orientation is assessed primarily within the imaging plane. Nevertheless, the workflow is modular and readily adaptable. Individual steps, including segmentation, model fitting, and orientation analysis can be replaced, and extensions to fully 3D fiber tracking is feasible when contributions from out-of-plane fibers are non-negligible. Overall, this framework provides a robust and adaptable approach for translating multiphoton microscopy images into quantitative descriptors of aortic wall microstructure relevant to studies of vascular remodeling.

## 5. Acknowledgments

The authors thank the Institute for Chemical Imaging of Living Systems (RRID:SCR 022681) at Northeastern University for consultation and instrument support. The authors also thank Dr. Cristina Cavinato (Université de Montpellier) for guidance on optimizing multiphoton imaging and interpreting SHG signals.

## 6. Funding Data

- National Institutes of Health (NIH) (Grant No. 1R01HL168473-01 to C.B.).
- National Science Foundation (NSF) (CAREER Award No. 2049088 to R.A.).
- American Heart Association (AHA) (Predoctoral Fellowship No. 24PRE1195859 to A.I.V.).

## 8. Data Availability Statement

The data generated and analyzed in this study are available from the corresponding authors upon reasonable request. The associated code is provided in the “Multiphoton Quantitative Analysis” repository on the Amini Lab Github website (AminiLabNU) (https://github.com/AminiLabNU/Quantitative-Analysis) [90].

## Abbreviations

2D: Two-dimensional
3D: Three-dimensional
CT-FIRE Curvelet Transform: Fiber Extraction
ECM: Extracellular Matrix
F1: F1 score
GT: Ground truth
IACUC: Institutional Animal Care and Use Committee
MPM: Multiphoton Microscopy
NAR: Nuclear Aspect Ratio
PCA: Principal Component Analysis
ROI: Region of Interest
SD: Standard deviation
SHG: Second Harmonic Generation (collagen channel)
TPEF: Two-photon Excited Fluorescence (fluorescent dyes)
TPEF-AF: Two-photon Autofluorescence (elastin channel)
VSMC: Vascular Smooth Muscle Cell

## 10. Nomenclature

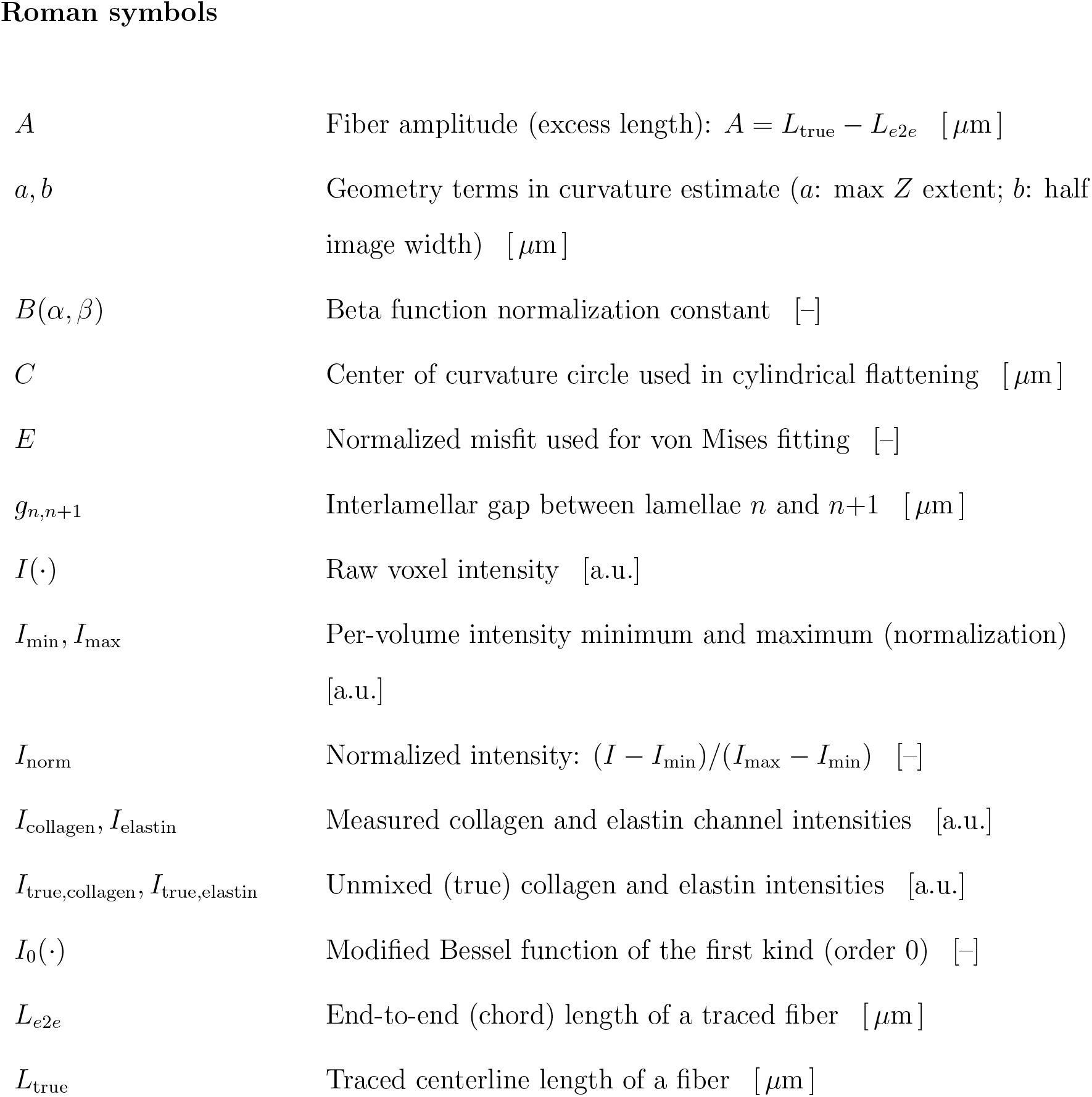

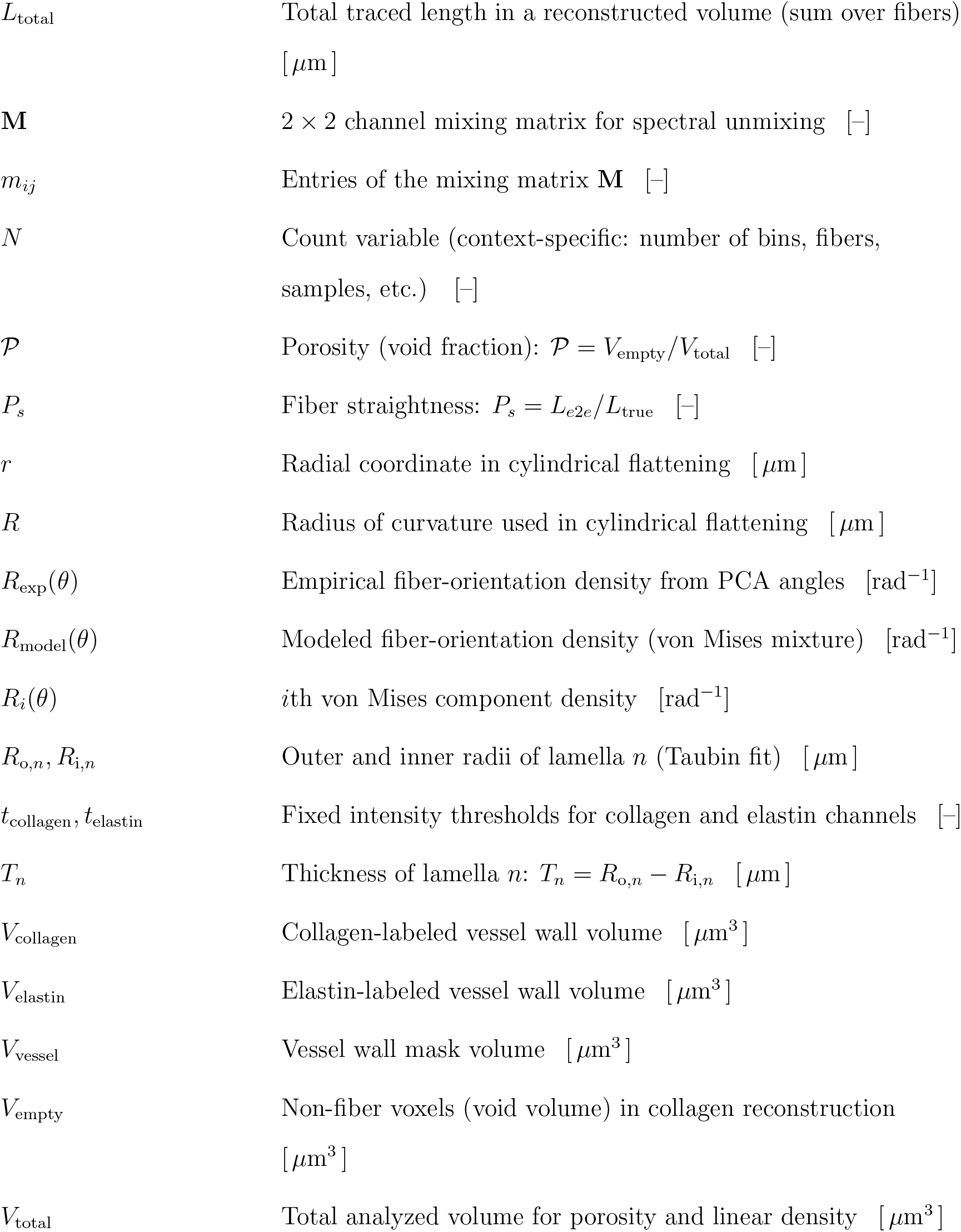

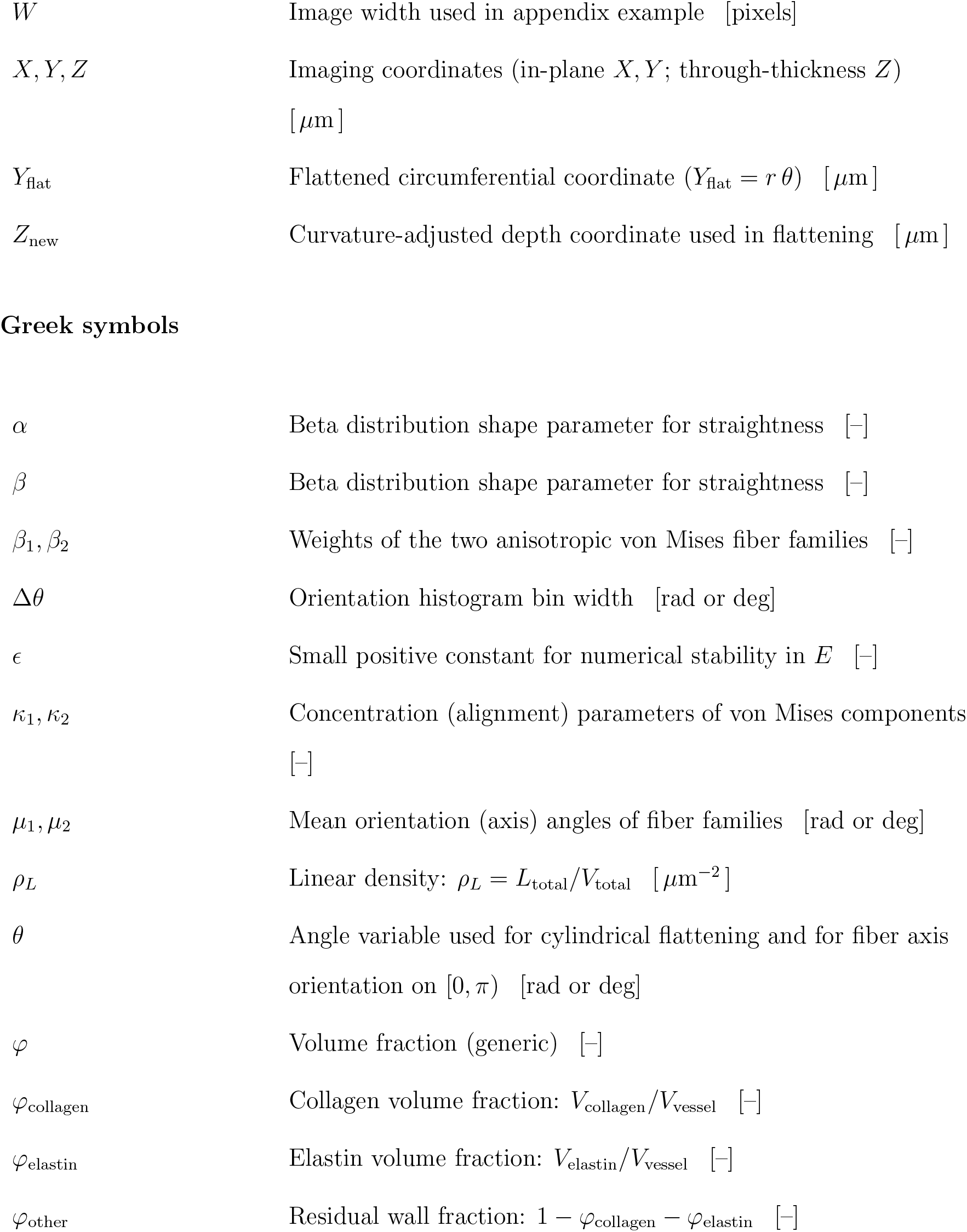

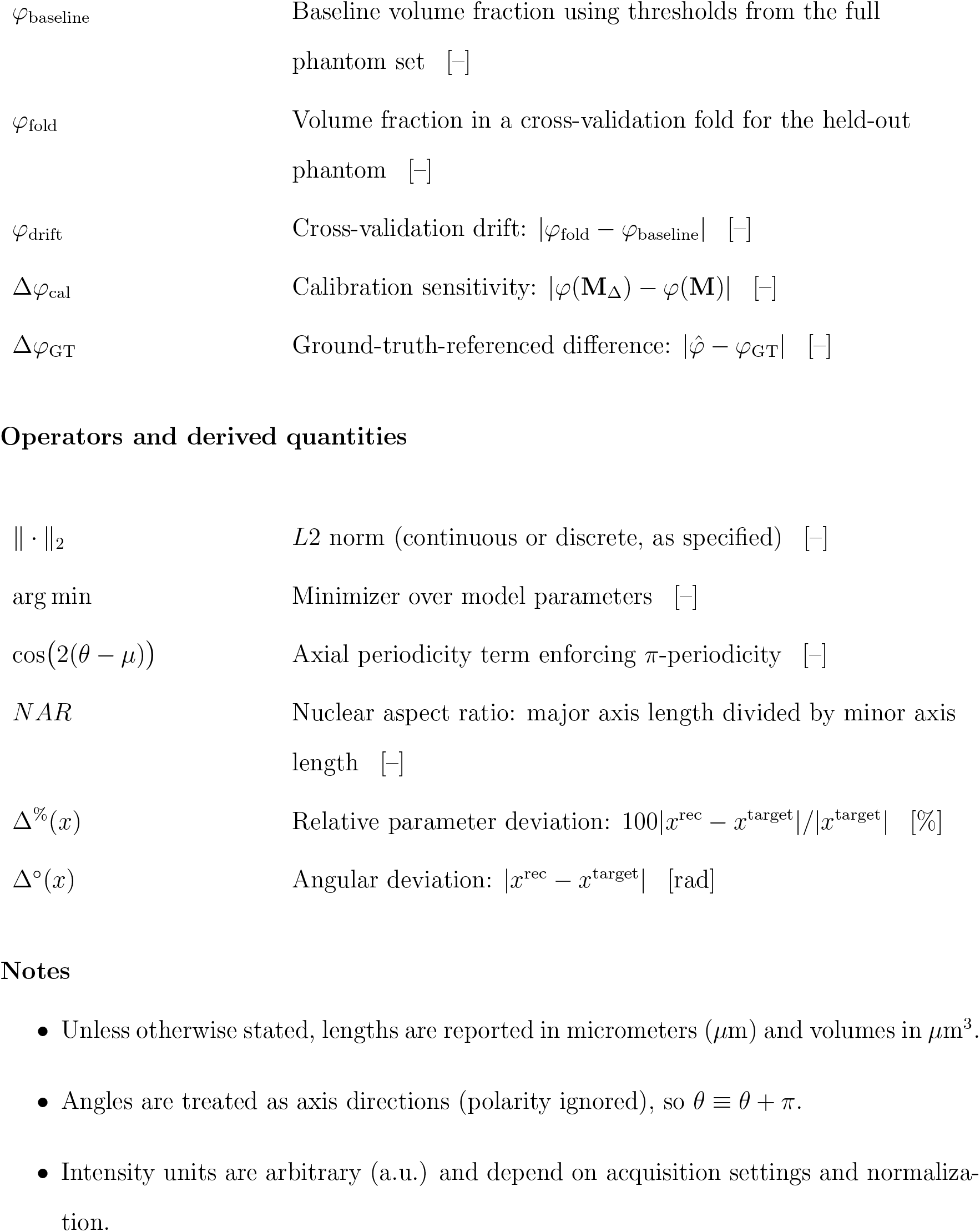

## Appendix

### Educational Materials: Cylindrical Flattening of Curved Vessel Walls

As in our prior efforts to integrate educational components within methodological papers [82, 83, 91, 92, 93, 94, 43, 95, 96], we provide here a concise homework problem designed for undergraduate students. The objective is to illustrate the concept of curvature correction and cylindrical flattening in vascular image analysis. The assignment is intended to be solved on paper without the use of computational tools.

### Problem

Multiphoton microscopy often produces images of vessel walls that are curved in cross-section. To facilitate quantitative analysis, these images can be transformed into a flattened coordinate system. Consider a *Y Z* image with pixel size 0.50 *μ*m/pixel, image width *W* = 400 pixels, and maximum axial index *z*_max_ = 80 pixels. The radius of curvature *R* is estimated as

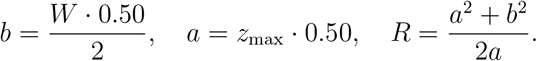

(Refer to section 2.4). For two points in the image, *P*_1_ : (*Y, Z*) = (60, 10) and *P*_2_ : (100, −5) (pixel coordinates), the cylindrical flattening transformation is defined by

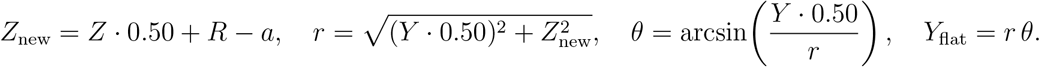

1. Compute *R* (in *μ*m).
2. For *P*_1_ and *P*_2_, calculate *Z*_new_, *r, θ* (both in radians and degrees), and *Y*_flat_ (all in *μ*m).
3. Determine which point lies farther along the circumferential direction in the flattened view.

